# Mitochondrial dysfunction impairs human neuronal development and reduces neuronal network activity and synchronicity

**DOI:** 10.1101/720227

**Authors:** T.M. Klein Gunnewiek, E. J. H. Van Hugte, M. Frega, G. Solé Guardia, K.B. Foreman, D. Panneman, B. Mossink, K. Linda, J.M. Keller, D. Schubert, D. Cassiman, E. Morava, R. Rodenburg, E. Perales-Clemente, T.J. Nelson, N. Nadif Kasri, T. Kozicz

**Affiliations:** Department of Anatomy, Radboudumc, Donders Institute for Brain, Cognition, and Behaviour, 6500 HB Nijmegen, the Netherlands; Department of Human Genetics, Radboudumc, Donders Institute for Brain, Cognition, and Behaviour, 6500 HB Nijmegen, the Netherlands; Department of Cognitive Neuroscience, Radboudumc, Donders Institute for Brain, Cognition and Behaviour, 6500 HB Nijmegen, the Netherlands; Department of Hepatology, UZ Leuven, 3000 Leuven, Belgium; Department of Clinical Genomics, Mayo Clinic, 55905 Rochester, MN, USA; Radboud Center for Mitochondrial Disorders, Radboudumc, 6500 HB Nijmegen, the Netherlands; Department of Laboratory Medicine and Pathology. Mayo Clinic, Rochester, MN 55905, USA; Departments of Medicine, Molecular Pharmacology and Experimental Therapeutics, Division of General Internal Medicine, Division of Paediatric Cardiology. Mayo Clinic Center for Regenerative Medicine, Rochester, MN 55905, USA; Department of Biochemistry and Molecular Biology, Mayo Clinic, 55905 Rochester, MN, USA

**Keywords:** MELAS, mitochondrial disease, mitochondria, neuron, induced pluripotent stem cells, network activity, neurodevelopment, micro-electrode array

## Abstract

Epilepsy, intellectual and cortical sensory deficits and psychiatric manifestations are among the most frequent manifestations of mitochondrial diseases. Yet, how mitochondrial dysfunction affects neural structure and function remains largely elusive. This is mostly due to the lack of a proper *in vitro* translational neuronal model system(s) with impaired energy metabolism. Leveraging the induced pluripotent stem cell technology, from a cohort of patients with the common pathogenic m.3243A>G variant of mitochondrial encephalomyopathy, lactic acidosis and stroke-like episodes (MELAS), we differentiated excitatory cortical neurons (iNeurons) with normal (low heteroplasmy) and impaired (high heteroplasmy) mitochondrial function on an isogenic nuclear DNA background. iNeurons with high levels of heteroplasmy exhibited mitochondrial dysfunction, delayed neural maturation, reduced dendritic complexity and fewer functional excitatory synapses. Micro-electrode array recordings of neuronal networks with high heteroplasmy displayed reduced network activity and decreased synchronous network bursting. The impaired neural energy metabolism of iNeurons compromising the structural and functional integrity of neurons and neural networks, could be the primary driver of increased susceptibility to neuropsychiatric manifestations of mitochondrial disease.

## Introduction

Mitochondrial disease is caused by mutations either in the nuclear or mitochondrial DNA (mtDNA). The resulting cellular/tissue energy crisis impacts organs with the highest energy need, such as the brain (El-Hattab et al., 2015). Epilepsy, intellectual and cortical sensory deficits and psychiatric manifestations are the most frequent manifestations of any mitochondrial disease (Srivastava et al., 2018; Andreazza et al., 2018; Sullivan et al., 2018; Sylvia et al., 2018; Kim et al., 2019; Finsterer, 2009; Gorman et al., 2016; Pei and Wallace, 2018; Reinhart and Nguyen, 2019). Neural processes with high energy demand, such as neuronal maturation/development and plasticity as well as impaired synaptic physiology and synchronous neuronal activity (Serafini, 2012; Alves et al., 2014; Boku et al., 2018; Quinn et al., 2018; Reinhart and Nguyen, 2019) could explain the proximal neuropsychological presentation in mitochondrial disease. To date, however, the lack of translational model systems of impaired energy flow in the brain has hampered our understanding of the exact nature of disease pathobiology and the development of disease-modifying therapies with a causal approach. Therefore, neural model systems are desperately needed.

**M**itochondrial **e**ncephalomyopathy, **l**actic **a**cidosis and **s**troke-like episodes *(MELAS)* is the most common progressive and devastating multisystem mitochondrial disease with epilepsy, stroke-like episodes, intellectual and cortical sensory deficits, psychopathology, muscle weakness, cardiomyopathy, and /or diabetes (El-Hattab et al., 2015). The majority of MELAS patients (80%) have an adenine to guanine pathogenic variant at the m.3243 position (m3243A>G) of the mitochondrial genome (mtDNA), in the *MT-TL1* gene specifying tRNA^leu(UUR)^ (OMIM 590050) (Goto et al., 1990; Hirano and Pavlakis, 1994). The estimated prevalence of clinically affected individuals with the m.3243A>G variant causing MELAS is about 1:20000 (Majamaa et al., 1998; Manwaring et al., 2007; Chinnery et al., 2000; Hirano and Pavlakis, 1994), but that of asymptomatic carriers could be as much as 1:400 in the general population (Manwaring et al., 2007). The percentage of mutated copies of mitochondrial DNA (heteroplasmy) plays a role in the onset and expression of symptoms, as well as the severity of the disease (Schon et al., 2012). Specifically, levels of heteroplasmy for the m.3243A>G variant positively correlates with mitochondrial respiratory chain complex I, III and IV insufficiencies (Ylikallio and Suomalainen, 2012; Ciafaloni et al., 1992; Yokota et al., 2015; Kobayashi et al., 1990, 1991). The m.3243A>G variant-related phenotypes are highly variable (Hamalainen et al., 2013). Lower heteroplasmy levels around 20% to 30% commonly present with type I or II diabetes, with or without hearing loss, while 50% to 80% can present with myopathy, cardiomyopathy, MELAS (Wallace, 2018). Homoplasmic cases are associated with either severe early onset MELAS, with encephalopathy including intellectual and cortical sensory deficits, psychiatric symptoms.

The polyploid nature of mtDNA and replicative segregation hamper the development of animal or *in vitro* disease models for mtDNA-related mitochondrial diseases (Prigione, 2015). Current *in vitro* models, such as cytoplasmic hybrids, do not take into account the interplay between patient mtDNA and nuclear DNA (Wilkins et al., 2014), important in both health (Latorre-Pellicer et al., 2016) and disease (Miller et al., 2011).

Human iPS cell are powerful tools in disease modelling and drug discovery through investigating the relationship between impaired brain energy metabolism in disease-relevant tissues and cell types, as well as to uncover aspects of the dynamic changes in neural structure and function that predispose to neuropsychological symptoms in mitochondrial disease (Srivastava et al., 2018). Using directed differentiation (Zhang et al., 2013; Frega et al., 2017), we generated human isogenic excitatory cortical neurons (iNeurons) containing low (0%) and high levels (>80%) of m.3243A>G heteroplasmy from patient-derived fibroblasts. We found that iNeurons with mitochondrial dysfunction exhibited reduced size and complexity of dendritic arbors, fewer excitatory synapses and accordingly a lower frequency of spontaneous postsynaptic currents. Further, neuronal network recordings from micro-electrode arrays showed less activity, as well as impaired synchronous network bursts of iNeurons with mitochondrial dysfunction. Our results highlight the potential of using iPS-derived neurons for disease modelling (Ben-Shachar and Ene, 2018), and provide conceptual advance in understanding the impact of mitochondrial dysfunction on the structural and functional integrity of neurons and neural networks.

## Results

### M.3243A>G heteroplasmy levels upon reprogramming and neuronal differentiation

We reprogrammed fibroblasts of an individual with MELAS to iPS cells by retroviral transduction with the Yamanaka transcription factors (see Supplemental data for patient description). We selected 6 clones and performed Sanger sequencing to determine the m.3243A>G heteroplasmy levels, and subsequently selected clones with low heteroplasmy (LH) (0% m.3243A>G) and with high heteroplasmy (HH) (71%- and 83% m.3243A>G), for pluripotency. Clones with and without m.3243A>G heteroplasmy showed positive expression of pluripotency markers OCT4, NANOG, SOX2, and LIN28, both via immunohistochemical staining and quantitative RT-PCR (Fig. S1A-D). We included a curated healthy iPS cell line (409-B, Kyoto University) in our study to serve as an external control. Next, we differentiated the iPS cells into a homogeneous population of excitatory cortical Layer 2/3 neurons (hereafter referred to as “iNeurons”) by forced expression of the transcription factor Ngn2 (Zhang et al., 2013; Frega et al., 2017). For all experiments, iNeurons were co-cultured on freshly isolated rodent astrocytes to facilitate neuronal maturation (Fig. S1F) (Frega et al., 2017). All iPS cell lines were able to differentiate into MAP2-positive excitatory iNeurons, which formed Synapsin 1/2 expressing synapses, within 21 days *in vitro* (DIV) of the start of differentiation (Fig. 1A; S1F-H). We noticed that differentiation induced higher mortality in the HH lines, which was estimated by the final number of surviving MAP2 positive iNeurons (Fig. S1G+H). We circumvented this issue by doubling the initial iPS cell plating density of the HH lines, whilst reducing the number of CTR- and LH cells, resulting in cultures with matching neuronal cell densities at DIV 21 (Fig. S1G+H, Materials and Methods). Importantly, high heteroplasmy level was retained across multiple passages of iPS cells (>60%, at least up to passage 40) and post differentiation into iNeurons (Fig. 1B).

**Figure 1.**
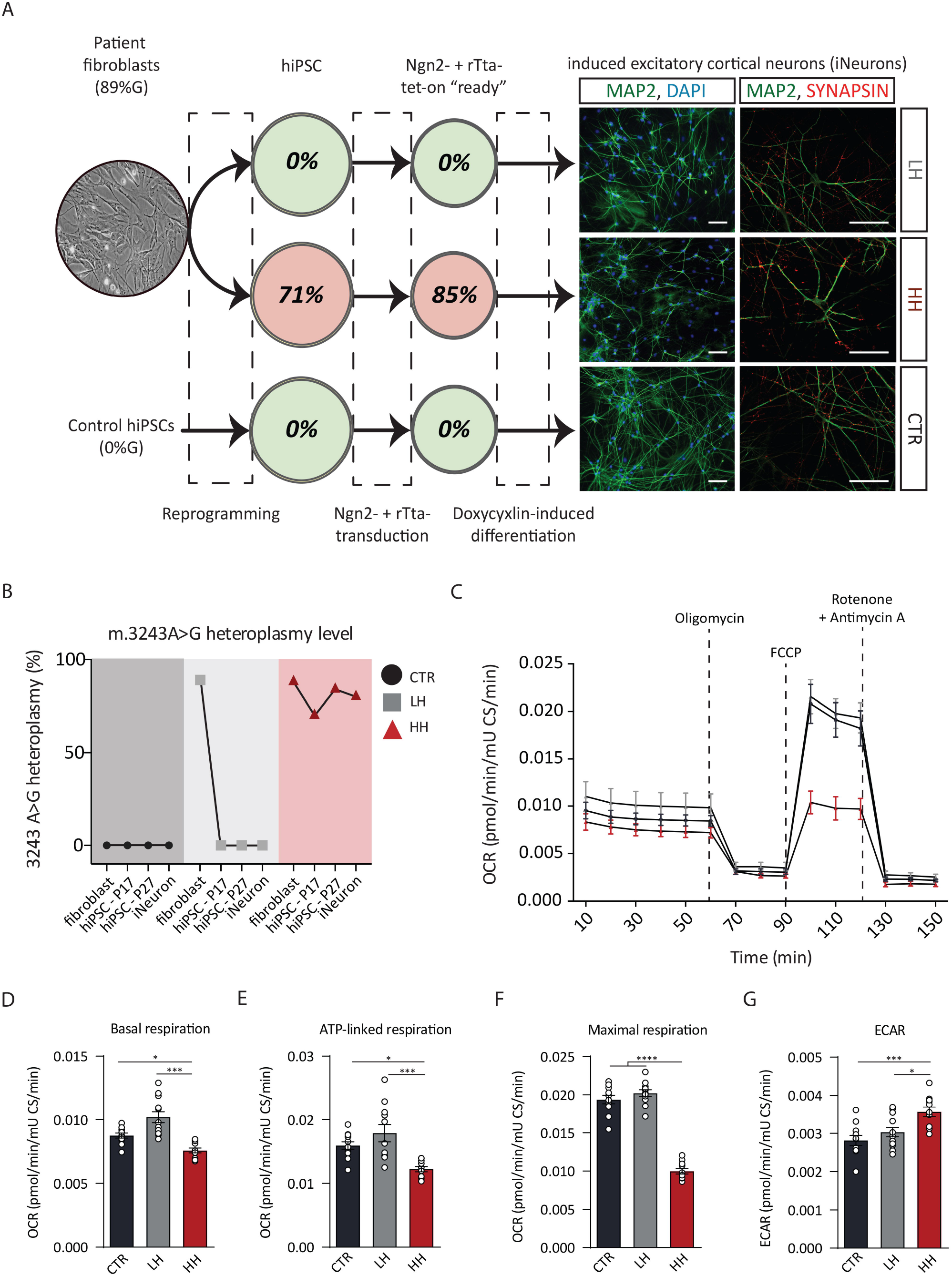
m.3243A>G heteroplasmy levels per cell type, and neuronal aerobic metabolic profiles. (A) Schematic representation of MELAS iPS cells and derived neurons. Patient fibroblast generated iPS clones with either homoplasmic (0%) or heteroplasmic (71%) mutation levels. Ngn2- and rTta-construct transduction led to 0% and 85% heteroplasmy levels. Subsequent doxycyclin-induced Ngn2-expression mediated differentiation into iNeurons, experssion micro-tubule associated protein 2 (MAP2) and Synapsin 1/2 at 21 days in vitro (DIV). (B) Sanger sequencing confirmed the biphasic mitochondrial segregation upon reprogramming of the fibrbolasts to iPS cells, as well as the retention of m.3243A>G heteroplasmy levels during during Ngn2-dependent differentiation. The color legend used here, is universal; dark grey for CTR, light grey for LH, and burgundy red for HH line(s). (C) Fifteen consecutive oxygen consumption rate (OCR) measurements, at basal level, and following supplementation with Oligomycin (2 μM), FCCP (2 μM), and Rotenone and Antimycin A (0.5 μM). Raw OCR levels were normalised to a citrate synthase assay (Fig. S2A). (D) Basal respiration was determined by averaging the first six recordings over all biological replicates (n = 12). (E) ATP-linked respiration (basal OCR minus Oligomycin response) was averaged over three measurements (n = 12). (F) Maximal respiration (FCCP response) was averaged over three measurements (n = 12). (G) Extracellular acidification rate (ECAR) (represents glycolysis rate) determined during- and averaged over-the first six recordings (n = 12); Raw ECAR (Fig. S2A) is normalised to a citrate synthase assay. Data represents means *±* SEM. *P<0.05, **P<0.01, ***P<0.001, ****P<0.001, one-way analysis of variance with post hoc Bonferroni correction. CTR-, LH- and HH iNeurons were statistically compared by One-way anova. C1, complex 1 (NADH dehydrogenase); C2, complex 2 (Succinate dehydrogenase); C3, complex 3 (Coenzyme Q: cytochrome c reductase); C4, complex 4 (Cytochrome c oxidase); C5, complex 5 (ATP synthase).

### High level of m.3243A>G heteroplasmy reduces mitochondrial oxygen consumption rate and mtDNA copy number

Neuronal differentiation induces a metabolic shift, from predominantly glycolytic iPS cells (Prigione et al., 2014) to mitochondrial oxidative phosphorylation (OXPHOS)-dependent neurons (Zheng et al., 2016). We assessed the effects of the m.3243A>G variant on mitochondrial respiration in control (409-B), LH and HH iNeurons with the Seahorse XF Cell Mito Stress Test. We used oxygen consumption rate (OCR) as a measure of mitochondrial respiration (Fig. 1C; Fig. S2A), extracellular acidification rate (ECAR) as a measure of glycolytic capacity (Fig. S2B), and we normalized the OCR/ECAR to oxaloacetate-induced citrate synthase activity (Rodenburg, 2011), to prevent a bias due to any differences in neuronal cell density (Fig. 1C; Fig. S2C). Control and LH iNeurons exhibited similar basal OCR profiles (Fig. 1C-G), whereas HH iNeurons showed a significantly lower basal OCR in comparison. Addition of the ATP-synthase inhibitor Oligomycin reduced ATP-dependent respiration (Fig. 1E) in HH iNeurons compared to both control and LH iNeurons. Furthermore, the uncoupling agent p-trifluoromethoxy carbonyl cyanide phenyl hydrazone (FCCP) showed a significantly reduced maximal respiratory capacity in HH iNeurons (Fig. 1F). Whereas OCR was decreased in the HH iNeurons under these multiple conditions, we observed an increase in ECAR in the HH iNeurons, reflecting an increase in anaerobic glycolysis and potential increase in lactate formation (Fig. 1G, Fig. S2). Overall, our data suggest that high m.3243A>G mutational load affects the OCR of mitochondria by reducing oxidative phosphorylation while increasing anaerobic glycolysis, and as a consequence reducing the efficiency of ATP production.

### Structural differences in iNeurons with high vs. low level of m.3243A>G heteroplasmy

Mitochondria support important aspects of neuronal development, such as axonal (Spillane et al., 2013) and dendritic branching (Agnihotri et al., 2017). We assessed the effects of m.3243A>G variant on somatodendritic neuronal structure by sparsely transfecting iNeurons at DIV 6 with a construct expressing red fluorescent protein (Fig. 2A). We imaged and reconstructed 3-dimensionally at least 30 iNeurons per cell line at DIV 23 (Fig. 2B) and quantified soma size, number of primary dendrites, total and mean dendritic length, number of dendritic nodes, and the surface covered by the dendritic trees (Fig. 2C). Control and LH iNeurons did not differ from each other in any of the parameters (Fig. 2C). Soma size and primary dendrite counts were similar between all cell lines, whereas we observed significantly shorter dendrites in HH iNeurons compared to control and LH iNeurons. Furthermore, we observed a significant reduction in total dendritic length, number of dendritic nodes and branching points in HH iNeurons. Accordingly, the total surface covered by the dendritic tree, quantified by calculating the convex hull, was significantly smaller in the HH iNeurons (Fig. 2C). Finally, Sholl analysis confirmed a reduced number of dendritic intersections, shorter dendritic length and fewer dendritic nodes per Sholl ring in HH iNeurons (Fig. 2D, E). These observations show that iNeurons with attenuated mitochondrial function are reduced and less complex dendritic organization and thus present a smaller receptive surface.

**Figure 2.**
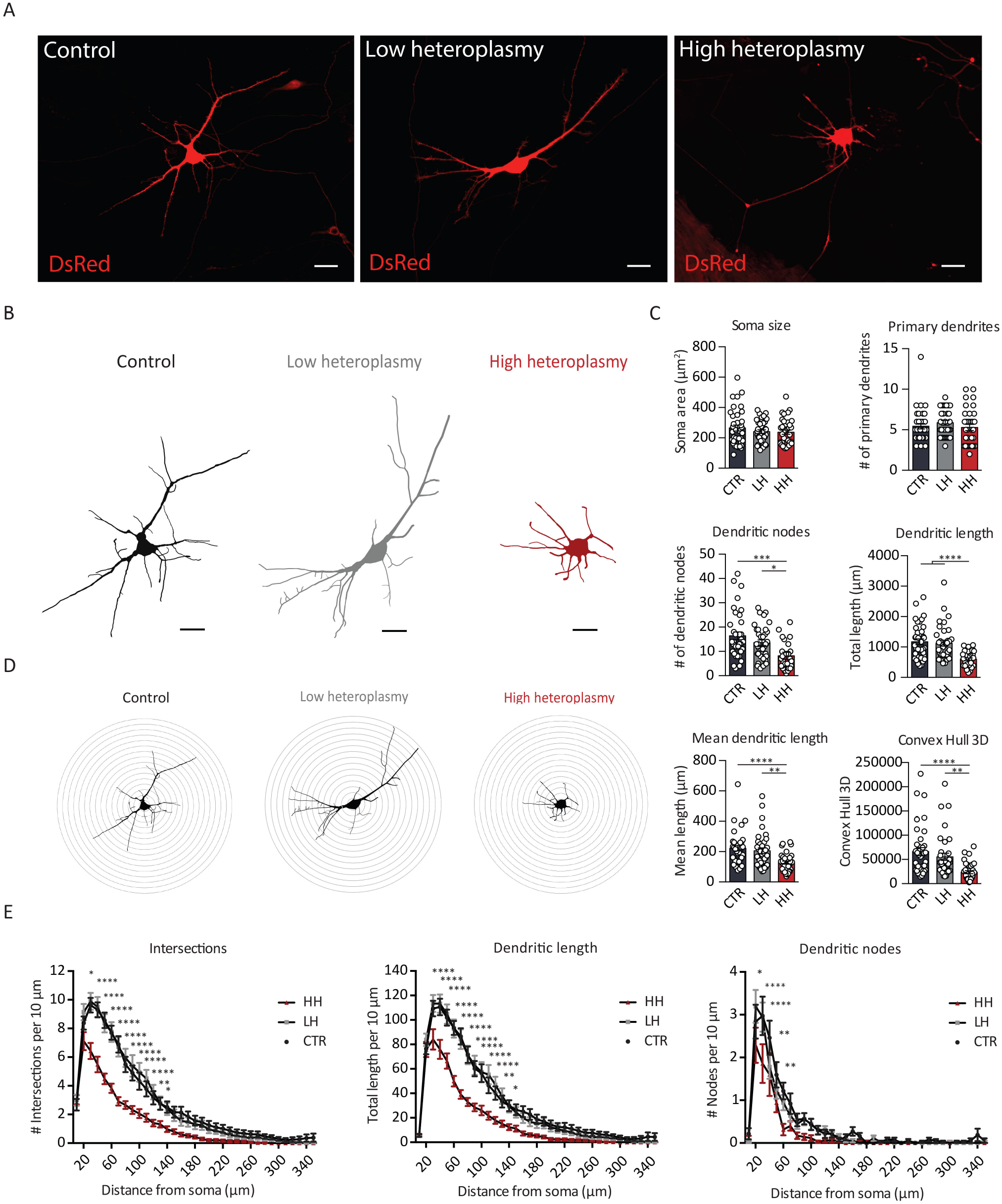
Reconstruction of the dendritic network of individual iNeurons. (A) Light fluorescence microscopy images of DsRed-positive iNeurons of CTR-, LH- and HH-cultures (scale bar = 30 μm). (B) Representative somatodendritic reconstructions of CTR-, LH- and HH iNeurons (CTR *n* = 37; LH *n* = 36; HH *n* = 35). C) Quantification of the soma size, the number of primary dendrites, the number of dendritic nodes, the mean- and total dendritic length, and the size of the surface covered by the dendritic network (Convex Hull 2D); *P<0.05; *** P<0.001, one-way analysis of variance. (D) Sequential 10 μm rings placed from the centre soma outwards for Sholl analysis. (E) Quantification per 10 μm Sholl section of the number of dendritic intersections per ring, the total dendritic length per ring, and the number of dendritic nodes per ring. Data represents means ± SEM. *P<0.05, **P<0.01, ***P<0.001, ****P<0.001, two-way analysis of variance with post hoc Bonferroni correction.

### Synaptic density and axonal mitochondrial abundance are reduced in iNeurons with high levels of m.3243A>G heteroplasmy

In addition to their role in neuronal growth, mitochondria mediate synapse formation and function, whether it is postsynaptic at dendrites (Li et al., 2004), at *en passant* pre-synaptic sites in the axon, or at growth cones (Morris and Hollenbeck, 1993; Smith and Gallo, 2018). Interestingly, mitochondrial absence at the synapse has been linked both to increased neurotransmitter release probability (Kwon et al., 2016) as well as to loss of synaptic function (Stowers et al., 2002). Our next goal was to investigate whether the m.3243A>G variant affects the number of synapses in iNeurons. To this end, we measured the number of synapses by quantifying presynaptic Synapsin-1/2 puncta, on MAP2-positive dendrites. Whilst we found no differences between control- and LH iNeurons, we did observe less Synapsin-1/2 puncta in the HH iNeurons compared to either the control or LH iNeurons (Fig. 3A).

**Figure 3.**
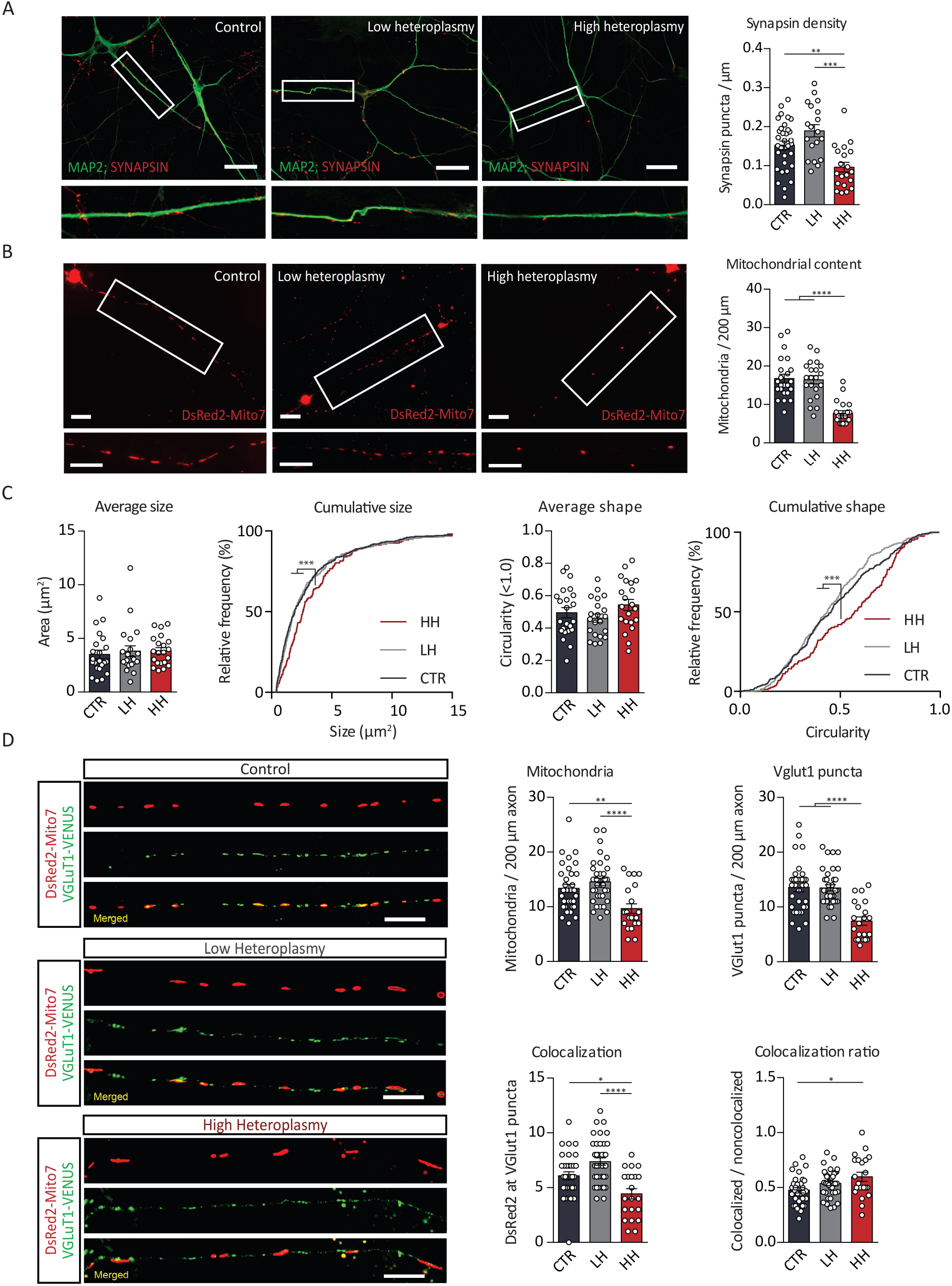
Mitochondrial- and synaptic density and co-localization. (A) Representative light fluorescence images of CTR-, LH-, and HH iNeurons (63X magnification) stained with micro-tubule associated protein 2 (MAP2; 488nm; dendritic marker) and Synapsin-1/2 (568nm; pre-synaptic protein) (Scale bar = 30μm) and quantification of the number of presynaptic Synapsin-1/2 puncta per μm MAP2-possitive dendritic surface, over a 100 μm section of a dendrite (CTR *n* = 21; LH *n* = 20; HH *n* = 20). Data represents means ± SEM. *P<0.05, **P<0.01, ***P<0.001, ****P<0.001, one-way analysis of variance with post hoc Bonferroni correction. (B) Light fluorescence images of CTR-, LH-, and HH iNeurons (40X magnification) transfected using DsRed2-Mito7 construct (Scale bar = 30 μm) as well as quantification of the number- and shape of mitochondrial particles in the initial proximal 200μm axon section (30μm from soma to exclude axon-initial segment) (CTR *n* = 23; LH *n* = 21; HH *n* = 21). (C) Average- and cumulative mitochondrial size and shape. ***P<0.001, Kolmogorov-Smirnov test. (D) Light fluorescence images of CTR-, LH-, and HH iNeurons co-transfected with a DsRed2-Mito7 (mitochondria) and a VGlut1-VENUS (VGlut1 puncta) constructs (Scale bar = 30 μm) and quantification of the number of mitochondria, the number of VGLut1 puncta, the absolute number of co-localizing (mitochondria plus VGlut1 puncta) and the ratio of co-localisation (co-localizing / non co-localizing) (CTR *n* = 23; LH *n* = 27; HH *n* = 21). Data represents means ± SEM. *P<0.05, **P<0.01, ***P<0.001, ****P<0.001, one-way analysis of variance with post hoc Bonferroni correction.

Next, we quantified the axonal mitochondrial abundance in the proximal part of the axon (30-200 μm from the soma). Using a DsRed2-Mito7 marker, we visualized the entire mitochondrial network of single iNeurons and found that HH iNeurons had fewer numbers of mitochondria present in the initial part of the axon (30-200μm from the soma) (Fig. 3B). Taking a more detailed look at the mitochondria, we found no differences in their average size, interconnectivity or shape (Fig. S3). However, cumulative distribution plots show that there was an increased proportion of larger and rounder mitochondria in HH iNeurons (Fig 3C).

Depending on species and neuronal subtype, mitochondria can accompany presynaptic sites (50% in human pyramidal neurons (Kwon et al., 2016); 82% in rat retinal ganglion neurons (Fischer et al., 2018); 43-56% in mouse hippocampal neurons (Obashi and Okabe, 2013)) where their presence can modify synaptic neurotransmitter release probability (Kwon et al., 2016; Werth and Thayer, 1994). To accurately determine the ratio of synapses co-localizing with mitochondria in human IPS-derived cortical neurons, we employed a dual DsRed2-Mito7 and VGLUT1-VENUS (pre-synaptic vesicular glutamate transporter) transfection. HH iNeurons again showed a reduced number of mitochondria in the distal axon compared to control and LH iNeurons (Fig. 3D), matching our observations in the proximal axon (Fig. 3A). Second, we observed a reduction in VGLUT1 puncta in HH iNeurons compared to control and LH iNeurons (Fig. 3D). The absolute number of VGLUT1 puncta that co-localized with mitochondria in the axon was lower in HH iNeurons than control and LH iNeurons, albeit the ratio of co-localizing vs. non-co-localizing synapses with mitochondria was comparable to both control and LH iNeurons. Similarly, no change in this ratio (synapses with versus synapses without mitochondria) was observed when using a DsRed2-Mito7 transfection in combination with an endogenous Synapsin staining, where we again confirmed the decreased number of synapses and axonal mitochondria in HH iNeurons (Fig. S10). This indicates that although the total number of synapses and mitochondria are reduced in HH iNeurons, no compensatory mechanism restores the absolute number of synapses co-localizing with mitochondria to levels observed in LH and control iNeurons.

In summary, we observed reduced numbers of mitochondria in the proximal and distal compartments of the axon in HH iNeurons, combined with fewer synapses. While the absolute number of synapses that contain mitochondria is reduced, the ratio of synapses that co-localize with mitochondria versus those that do not is stable across all cell lines.

### Frequency of spontaneous excitatory activity is reduced in iNeurons with high level of m.3243A>G heteroplasmy

After finding a deficit in mitochondrial function, as well as a decrease in neuronal complexity, synaptic density, and mitochondrial abundance, we next sought to study the effects of m.3243A>G heteroplasmy on neuronal activity at the single-cell level, using whole cell voltage-clamp recordings. We recorded spontaneous excitatory post-synaptic currents (sEPSCs) at −60 mV, for all three neuronal lines (Fig 4A). We observed a decrease in frequency, but not amplitude, of sEPSCs in the HH iNeurons, as compared to the control and LH iNeurons (Fig 4B).

**Figure 4.**
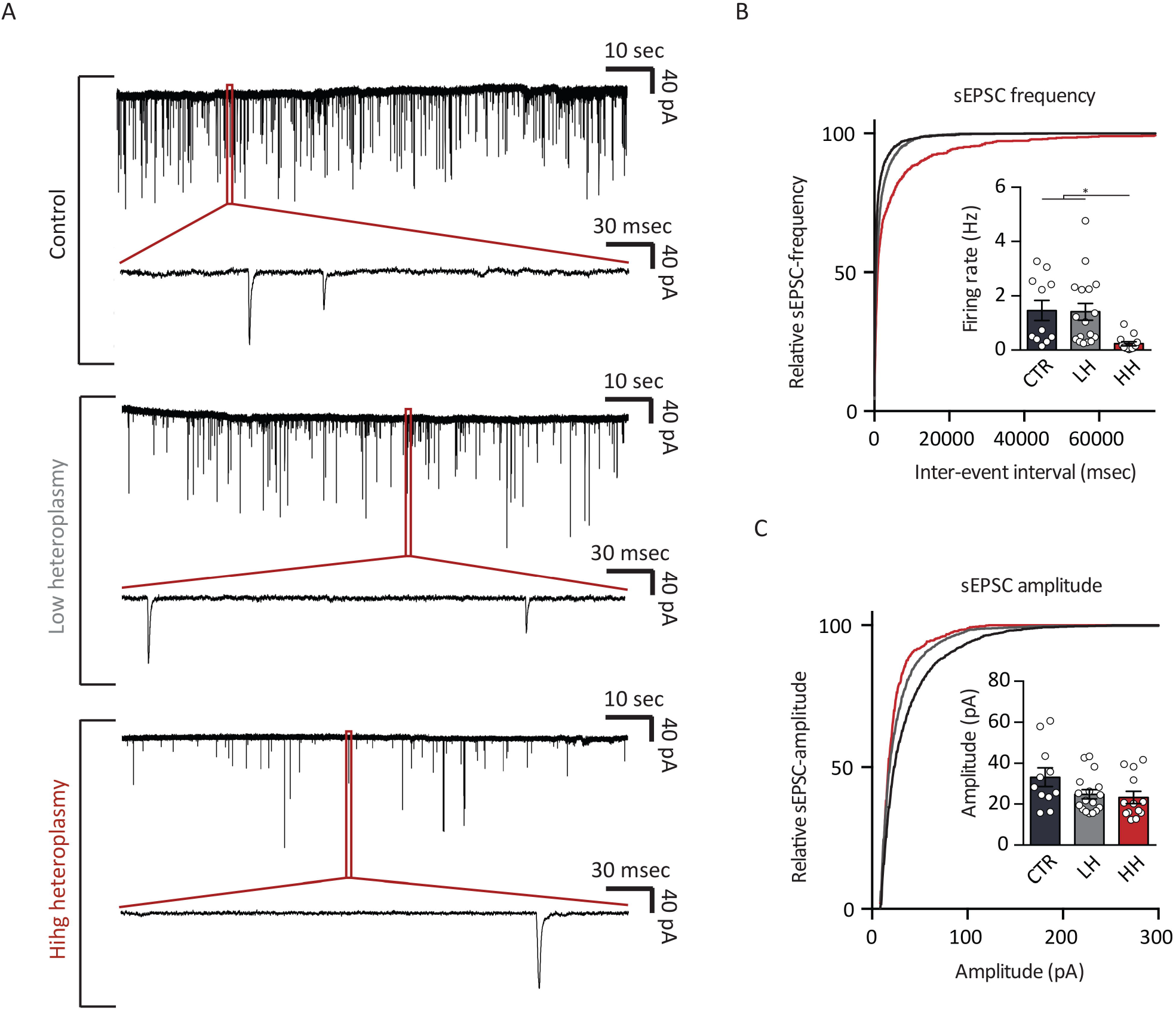
Spontaneous excitatory activity at the single-cell level. (A) Representative electrophysiological traces showing spontaneous excitatory post-synaptic currents (sEPSCs) recorded at −60mV, using Voltage Clamp whole-cell recording in iNeuron cultures at DIV 23 (CTR *n* = 12; LH *n* = 17; HH *n* = 13). (B) sEPSC frequency and (C) sEPSC amplitude were compared between CTR-, LH-, and HH iNeurons cultures. Data represents means ± SEM. *P<0.05, **P<0.01, ***P<0.001, ****P<0.001, one-way analysis of variance with post hoc Bonferroni correction.

This m.3243A>G variant-induced reduction in sEPSC frequency is in accordance with our aforementioned reduction in synaptic density. Taken together, we observed correlates of impaired neuronal energy metabolism not only on neuronal morphology, but also on single neuronal function.

### High level of m.3243A>G heteroplasmy impairs neuronal network activity and synchronicity

The previous experiments showed that, at the single cell level, iNeurons with high m.3243A>G heteroplasmy form fewer synapses and receive less synaptic input. We next investigated whether this reduced synaptic activity also translated into altered activity at the network level. To this end we examined and compared the spontaneous activity of neuronal networks derived from LH and HH iNeurons growing on micro-electrode arrays (MEAs) at similar densities (Fig. S4A). MEA recordings allowed us to non-invasively and repeatedly monitor neuronal network activity (spikes and bursts) through extracellular electrodes located at spatially separated points across iNeuron cultures (Fig. 5A-C). Because of the known heterogeneity seen between MELAS patients carrying the m.3243A>G heteroplasmy, and to avoid a potential mediating effect of the patient’s specific genetic background, we generated three additional sets of isogenic MELAS iPS lines (LH2-4 and HH2-4). We successfully selected homoplasmic clones with low heteroplasmy (LH2-4; 0% m.3243A>G) and with high heteroplasmy (HH2-4; 84%-m.3243A>G).

**Figure 5.**
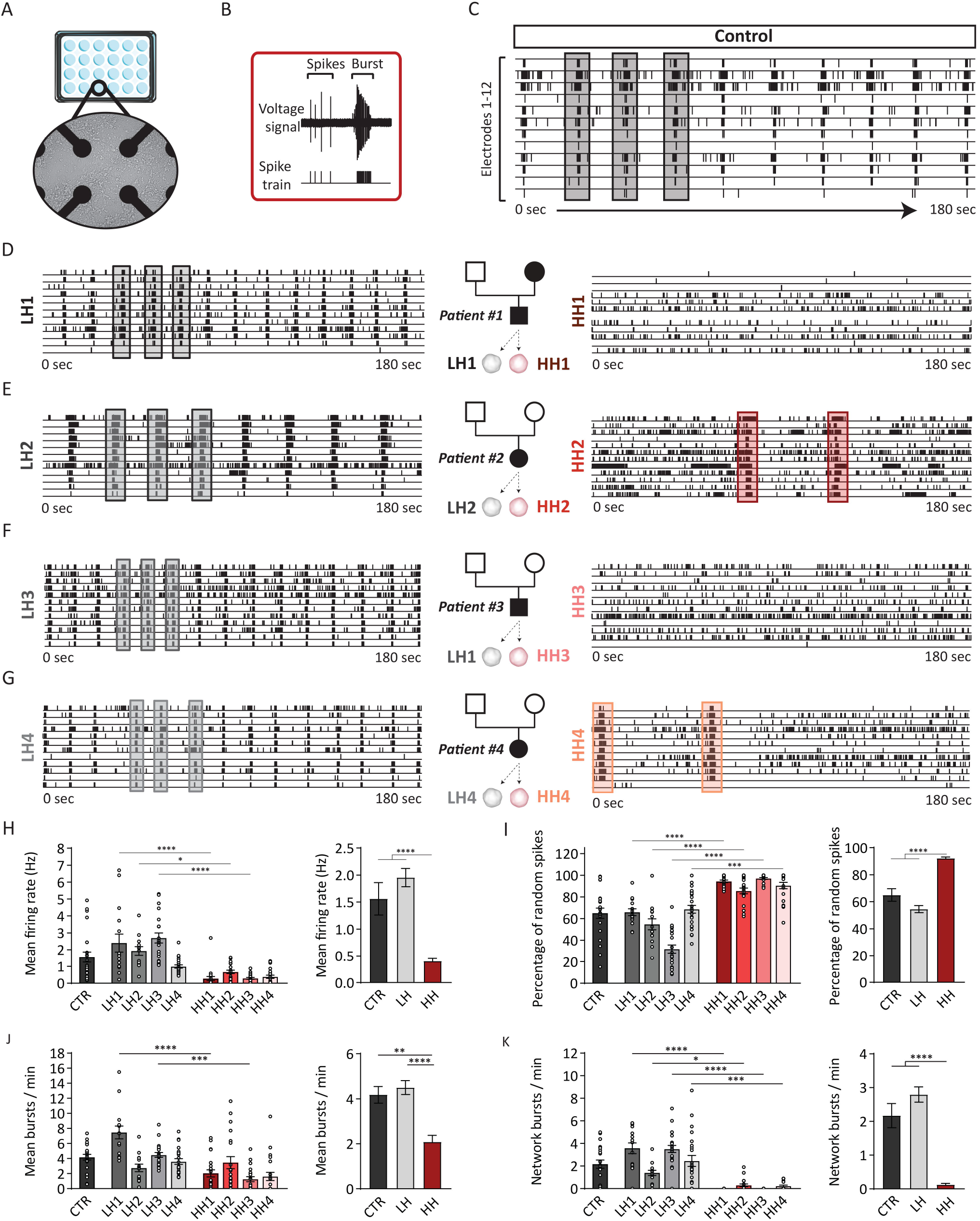
Spontaneous excitatory network activity recorded on MEA. (A) CTR, LH1-4, and HH1-4 neuronal networks cultured on 24-well micro-electrode arrays (MEA) (CTR n = 23; LH1 n = 15; LH2 = 14; LH3 = 22; LH4 = 23; HH1 n = 22; HH2 = 20; HH3 = 21; HH4 = 21); and recorded for spontaneous excitatory activity at DIV30. The iNeurons are shown here in, grown in high density, on top of the microelectrodes. (B) Spikes and bursts from a 180-second recording plotted per electrode, (C) CTR-network plotted here as example, showing synchronous network bursts over all electrodes (grey vertical bar), over a 3-minute period. (D) Example plots of LH1- and HH1 networks derived from patient #1. (E) Example plots of LH2- and HH2 networks derived from patient #2. (F) Example plots of LH3- and HH3 networks derived from patient #3. (G) Example plots of LH4- and HH4 networks derived from patient #4. We quantified (H) mean firing rate (MFR), (I) percentage of random spikes (PRS), (J) mean burst rate, and (K) network burst rate (per minute; NBR). Data represents means ± SEM. *P<0.05, **P<0.01, ***P<0.001, ****P<0.001, one-way analysis of variance with post hoc Bonferroni correction.

During the fifth week in vitro (DIV 30), all control neuronal networks (i.e. CTR, LH1, LH2, LH3 and LH4) showed a pattern of activity characterized by regular synchronous events, called network bursts (Fig. 5C-G). These network bursts are an important characteristic of a properly developed, mature neuronal network (Frega et al., 2017). At this stage, we observed no difference in the level and pattern of synchronous activity between the CTR, LH1-4 neuronal networks (Fig 5C-K). The highly reproducible network characteristics observed across all controls and LH1-4 lines provided us with a consistent and robust standard to which we could directly compare the HH1-4 networks.

The iNeurons with high levels of m.3243A>G heteroplasmy (HH1-4) showed spontaneous activity with bursts (Fig. 5D-G) of a relatively similar duration (Fig. S4B) and comparable number of spikes as iNeurons with LH (Fig. S4C, burst firing rate). However, both the amount and pattern of spike and network bursting in HH1-4 neuronal networks were significantly different compared to their respective LH isogenic controls. In particular, we found that the general level of activity (mean firing rate; MFR) exhibited by the HH1-4 neuronal networks was strongly reduced (Fig. 5H). Furthermore, HH1-4 neuronal networks exhibited a strong reduction in the network burst rate (NBR) (Fig. 5K), with HH1 and HH3 networks exhibiting virtually no network bursts (Fig. 5D, F, K). Network burst duration (NBD) in HH2 and HH4 networks, however, were not affected (Fig. S4B). Notably, because network bursts were very sparse in HH1 and HH3 networks, we were unable to calculate the NBD for these. Finally, we also observed that spike organization in HH1-4 neuronal networks differed from controls, which was indicated by the higher percentage of random spikes (PRS) occurring outside the network bursts (Fig. 5I). Taken together, these results show that iNeurons with high levels of m.3243A>G heteroplasmy fail to organize into conventional functional neuronal networks and produce a distinctive phenotypical pattern of network activity.

Next, we assessed if and to what degree control and LH network differed from HH neuronal networks when taking all measured network parameters into account. To this end we performed a canonical discriminant function analysis including MFR (Fig. 5H), PRS (Fig. 5I), Mean burst rate (MBR, Fig 5J), Mean burst duration (MBD, Fig. S4B), Mean burst firing rate (MBFR, Fig. S4C), Mean burst interval (MBI, Fig. S4D), NBR (Fig. 5K), NBD (Fig. S4E), and the network inter-burst interval (NIBI, Fig. S4F) as variables (Fig. S4G). The analysis revealed that HH groups clearly clustered outside the CTR/LH spectrum (Fig. S4G). Based on this model, we could not only predict an individual’s membership in the larger known group, but also cluster them into the respective subgroups (Fig. S4H). Combined, the data show that abnormal energy metabolism in iNeurons impacts neuronal network organization and activity, resulting in a distinct neuronal network phenotype.

## Discussion

Here we report on the development and deep phenotyping of a translational neural model system of mtDNA-related mitochondrial disease. Specifically, we used state-of-the-art iPS cell technology (multiple patient lines and leveraging on naturally occurring isogenic lines with various levels of mitochondrial dysfunction), combined with in depth morphological and electrophysioloigical phenotyping (network and single cell level) to identify a hitherto unknown cell- and network-level neural phenotypes of mitochondrial disease.

iPS cell reprogramming produced clones with different heteroplasmy levels, including homoplasmic clones, due to natural underlying heterogeneity in the original fibroblast population with concomitant changes in respiratory chain activity (this study and Perales-Clemente et al., 2016). This allowed us to select iNeurons with isogenic nuclear background (eliminating a potential confounding effect due to differences in nuclear genetic background) and with appropriate heteroplasmy levels and respiratory functions for disease modelling. We observed that iNeurons generated from individuals with the m.3243A>G variant faithfully replicate brain-specific manifestations of respiratory complex deficiency. iNeurons also revealed clues for understanding the pathomechanism related to abnormal energy metabolism in the brain. Specifically, we found evidence that iNeurons with high level of m.3243A>G heteroplasmy exhibited reduced dendritic length and complexity. Furthermore, we found a reduction in the number of mitochondria in the proximal section of the axon, combined with a reduction in pre-synaptic protein abundance. On a functional level, iNeurons with high levels of m.3243A>G heteroplasmy were less active, both at the single neuron and neuronal network level, and showed a reduction in network synchronicity.

Neuropathological studies have expanded our understanding of neural impairment and cell loss in the brains of individuals with mitochondrial disease revealing structural alterations. Mitochondrial dysfunction has been associated with altered neuronal dendritic morphology and remodelling (Tsuyama et al., 2017). The local availability of mitochondrial mass is critical for generating and sustaining dendritic arbors, and disruption of mitochondrial distribution in mature neurons is associated with structural alterations in the neurons (López-Doménech et al., 2016; Spillane et al., 2013; Kuzawa et al., 2014). Loss of interneurons (Lax et al., 2016), synapses and dendritic atrophy (both in number and size) in specific brain areas, such as various cortical areas, cerebellum, thalamus and basal ganglia (Cobley, 2018; Chen et al., 2017; Quintana et al., 2010; Betts et al., 2006; Turnbull et al., 2010; Briston and Hicks, 2018) have also been reported in individuals with mitochondrial dysfunction. We found a reduced number of primary dendrites and dendritic nodes, decreased total- and mean dendritic length, as well as reduced surface, covered by the dendritic network. We also observed reduced synaptic density in iNeurons with high levels of heteroplasmy concomitant with a reduction in the number of mitochondria in the proximal- and distal section of the axon. Our observations, therefore, corroborate the notion that mitochondrial function and positioning influencing dendritic branch morphology plays a direct role in establishing and maintaining mature neuronal circuits.

iNeurons with high levels of m.3243A.G heteroplasmy and grown on MEA exhibited strongly reduced mean firing rate (MFR). Neuronal activity, being heavily dependent on glucose supply as the main fuel source (Magistretti and Allaman, 2015), is especially vulnerable to metabolic dysregulation. Thus, pathological brain states, such as epilepsy characterized by firing instability and recurrent seizures, reflecting aberrant synchronous activity of large groups of neurons are likely to be associated with impaired energy flows in the brain. In a recent study, Styr *et al.,* (2019) observed that metabolic signalling constitutes a core regulatory module of MFR homeostasis. Our results confirm the link between neuronal energy metabolism and MFR homeostasis as well as corroborate the notion that mitochondrial dysfunctions could be causal in the initiation and progression of distinct types of epilepsy (Zsurka and Kunz, 2015).

Synchronous rhythms represent a core mechanism for sculpting temporal coordination of neural activity in brain-wide networks (Buzsáki et al., 2013). Common symptoms of m.3243A>G related mitochondrial disease, such as epilepsy, intellectual and cortical sensory deficits, as well as psychopathology are characterized too by excessive or asynchronous neural activity that typically does not occur in a single isolated neuron; rather, it results from pathological activity in large groups-or circuits-of neurons (Leistedt and Linkowski, 2013; Alexander et al., 2016; Beghi et al., 2005; Lenartowicz et al., 2018; Wang, 2010). Energy is required to fuel the formation and synchronized activity of neuronal circuits (Jan and Jan, 2010; Spruston, 2008). Altered energy metabolism in the brain, therefore, leads to dramatic changes in the underlying synchronization and functioning of these networks (Quinn et al., 2018; Styr et al., 2019). Impaired mitochondrial structure and function predispose neuronal network dysfunction (Virlogeux et al., 2018). Various *in vitro* and *in vivo* model systems have also shown impaired neuronal oscillatory function in mitochondrial disease models (for review see; Chan *et al.,* 2016). Our observation of a reduced level of neuronal network activity as well as the pattern of spiking and the bursting activity of neuronal networks caused by high heteroplasmy levels supports the notion that neurons with mitochondrial dysfunction fail to organize into functional neuronal networks. Regulation of synchronous brain activity in individuals with mitochondrial a disease therefore could be proximal to the clinical phenotype of epilepsy, cognitive impairment and neuropsychiatry.

Taken together, investigating the relationship between impaired brain energy metabolism in a disease-relevant tissue recapitulated similar structural and functional neuronal phenotypes that exist in epilepsy, intellectual and cortical sensory deficits. These results suggest that mitochondrial dysfunction could be a primary driver in the initiation and progression of neuropsychiatric symptoms in individuals with mitochondrial disease. These results go beyond of the etiology of MELAS disease and do provide conceptual advance in understanding the impact of mitochondrial dysfunction on the structure and function of single neurons and neural networks. Our results not only help understand the pathobiology of common neuropsychiatric symptoms in mitochondrial disease, but could be leveraged for future pharmacological studies, which are currently only performed on fibroblast due to lack of a proper mitochondrial disease model system.

## Acknowledgements

We thank EM and DC of the University Hospital Leuven, and EPC and TJN of the Mayo Clinic, for their generous donation of the patient derived IPS lines. We thank F. Polleux (Columbia University, NY) for sharing the VGLUT1-VENUS construct. This work was supported by the Tjalling Roorda Foundation (to TMKG), Stichting Stofwisselingskracht (to TK and NNK) and open ALW ALW2PJ/13082 (to NNK).

## Author contributions

TMKG, TK, and NNK conceived and supervised the study. TMKG, EJHVH, MF, GSG, BM, KBF, DP, KL, and JMK assisted in the performance and/or analysis of the experiments. DS, DC, EM, RR, NNK and TK provided facilities or equipment. DC, EM, EPC, and TJN provided either patient fibroblasts or patient derived IPS lines. TMKG, MF, TK, and NNK wrote the manuscript. All authors reviewed and edited the manuscript. TMKG, NNK, and TK carried out funding acquisition.

## Materials and Methods

### Subjects

Subject #1 was born, as a second child of a then healthy 29-year-old mother, who was later diagnosed with a classic MELAS phenotype, due to 3243A>G mutations. The older sister, who developed severe depression around the age of 32 years, and had recurrent episodes of visual loss, received the same diagnosis after her first stroke-like episode. Our patient had normal growth and adequate early psychomotor development. During puberty he was evaluated for exercise intolerance and fatigue. He was diagnosed with cardiomyopathy at the age of 31 years. He became obese and was diagnosed with type 2 diabetes mellitus. He developed severe, therapy-resistant depression at the age of 33 years. He was found to have homoplasmic 3243A>G mutations in urine sediment cells, 58% heteroplasmy in blood and 89% in fibroblasts. Further metabolic evaluation detected elevated lactic acid levels. Additional BAER test detected unilateral sensorineural hearing loss. At the age of 42 years our patient is using a wheelchair due to progressive muscle weakness and has a mild asymmetry in muscle strength after two stroke-like episodes. His cardiac disease is stable on beta-blocker therapy. His hearing loss became bilateral.

Subject #2 had no family history of mitochondrial disease. Her only notable clinical feature in the paediatric age was a childhood onset progressive sensorineural hearing loss, which needed a left cochlear implant treatment. During puberty she developed insulin dependent Diabetes mellitus and depression. She was also treated for chronic constipation. She had no other health concerns till the age of 36 years, when she had the first of his recurrent stroke like episodes. She also developed seizures and gait ataxia and has a slowly progressive cognitive decline in association with the recurrent stroke like episodes.

Subject #3 had no significant family history and had an uneventful early childhood. He was investigated for childhood onset recurrent vomiting. As a young adult he developed insulin dependent Diabetes mellitus. He also had his first episode of major depression at the age of 25 years. He was diagnosed with sensorineural hearing loss. At the age of 30 years he had the first of his recurrent stroke like episodes. He also developed recurrent complex migraine episodes with visual loss. He developed rapid cognitive decline in association with seizures and behavioural difficulties. In one year, he was not able to care for himself anymore. He was found to have heteroplasmic 3243A>G mutations in 86% in urine sediment cells, 28% heteroplasmy in blood and 30% in fibroblasts.

Subject #4 had a history of diabetes and deafness in his mother. Her paediatric disease history was uneventful. At the age of 23 years she developed insulin dependent Diabetes mellitus. At the age of 25 years she was diagnosed with hearing loss and had her first episode of major depression. She was also diagnosed with nephropathy and microalbuminuria at the age of 25 years. At the age of 30 years she had the first of her recurrent stroke like episodes. At the age of 35 years she was treated for concentration and memory loss and suicidal ideas. At age 37 she was diagnosed with ophthalmoplegia, ptosis and myopathy, developed chronic fatigue, exercise intolerance and in her late 30s peripheral neuropathy. Her MRI showed brain atrophy with subcortical white matter lesions and basal ganglia involvement. At age 45 she was found to have heteroplasmic 3243A>G mutations in 47% in urine sediment cells, 16% heteroplasmy in blood and 33% in fibroblasts.

### IPS cell generation and culture

Skin fibroblasts of MELAS subject #1 with the pathogenic variant m.3243A>G in *MT-TL1* (tRNA^Leu(UUR^ were reprogrammed by the in house Radboudumc Stem Cell Technology Centre, through retroviral transduction of the Yamanaka transcription factors Oct4, c-Myc, Sox2 and Klf4 (Takahashi and Yamanaka, 2006). IPS cells for subject #2-4 (see Supplemental data for patient description) were reprogrammed from primary human skin fibroblasts by ReGen Theranostics (Rochester, MN) using CytoTune-iPS 2.0 Sendai Reprogramming Kit according to manufacturer’s instructions (Invitrogen A16517). M.3243A>G heteroplasmy levels of iPS lines were sequenced, after which one iPS clone without-, and one iPS clone with heteroplasmy were selected. An additional, curated control cell line (409B2 (Kondo et al., 2017)) was included, and confirmed to have no m.3243A>G heteroplasmy. IPS cells were thawed and cultured at all times cultured on recombinant human laminin LN521 (Biolamina, 2021-21) kept in E8 basal medium (Thermo Fisher Scientific) supplemented with primocin (0.1 μg/ml, Invivogen), puromycin (0.5 μg/ml) and G418 (0.5 μg/ml), and uridine (50 μg/ml) and incubated at 37°C/5%CO2, with medium changes every 2-3 days and passages 1-2 times per week using ReLeSR (Stem Cell Technologies). Heteroplasmy levels were regularly measured to insure they were retained across passage numbers. All measurements were performed on iPS cells with passage lower than 40.

### Neuronal differentiation

IPS cells were directly derived into upper-layer, excitatory cortical neurons by overexpressing the neuronal determinant Neurogenin 2 (Ngn2) upon doxycycline treatment based on (Zhang et al., 2013), and as we described previously (Frega et al., 2017). RtTA/Ngn2-positive iPS cells were plated as single cells at DIV0 onto nitric-acid treated coverslips coated with 50μg/mL poly-L-ornithine hydrobromide (PLO; Sigma-Aldrich #P3655-10MG) and 5 μg/mL human recombinant laminin 521 (BioLamina #LN521-02) in E8 basal medium (Gibco #A1517001) supplemented with 1% Penicillin/Streptomycin (Pen/Strep; Sigma-Aldrich P4333), 1% RevitaCell (Thermo-Fisher #A2644501), and 4 μg/mL doxycycline (Sigma-Aldrich #D9891-5G) to drive Ngn2 expression, incubated at 37°C/5%CO2, at 20,000 cells per well (24 well plate) for the CTR- and LH-lines, and 30,000 cells per well (24 well plate) for HH lines. At DIV1, medium was changed to DMEM/F12 (Gibco #11320-074) supplemented with 1% Pen/Strep, 4 μg/mL doxycycline, 1% N-2 supplement (Gibco #17502-048), 1% MEM non-essential amino acid solution (NEAA; Sigma-Aldrich #M7145), 10 ng/mL human recombinant BDNF (Promokine #C66212), 10 ng/mL human recombinant NT-3 (Promokine #C66425). To support the neuronal culture, freshly prepared primary cortical rat astrocytes (Frega et al., 2017) were added on DIV2, in a 1:1 ratio. At DIV3, the medium was changed to Neurobasal (Gibco #21103-049), supplemented with 20 μg/mL B-27 (Gibco #0080085SA), 1% GlutaMAX (Gibco), 1% Pen/Strep, 4 μg/mL doxycycline, 10 ng/mL human recombinant NT3, 10 ng/mL human recombinant BDNF. At DIV3 only, 2 μM Cytosine b-D-arabinofuranoside (Ara-C; Sigma-Aldrich C1768-100MG) was added to remove any proliferating cells. From DIV5-DIV9 50% of the medium was refreshed every 2 days. From DIV 9-23 onwards the medium was supplemented with 2,5% Fetal Bovine Serum (FBS; Sigma-Aldrich #F2442-500ML) to support astrocyte viability.

### Seahorse Mito Stress Test

Oxygen consumption rates (OCR) were measured using the Seahorse XFe96 Extracellular Flux analyser (Seahorse Bioscience). IPS cells were seeded at a concentration of 10,000 per well in E8 basal medium supplemented with 1% Pen/Strep, 10 μg/mL RevitaCell, and 4 μg/mL doxycycline, and allowed to adhere at 37 °C and 5% CO_2_. A similar maturation protocol was applied as previously described, up until the cells were 23 days in vitro. One hour before measurement, culture medium was removed and replaced by Agilent Seahorse XF Base Medium (Agilent #103334-100) supplemented with 10 mM glucose (Sigma), 1 mM sodium pyruvate (Gibco), and 200 mM L-glutamine (Life sciences) and incubated at 37 °C without CO2. Basal oxygen consumption was measured six times followed by three measurements after each addition of 1 μM of oligomycin A (Sigma #75351-gMG), 2 μM carbonyl cyanide 4-(trifluoromethoxy) phenylhydrazone FCCP (Sigma #C2920-50MG), and 0. 5 μM of rotenone (Sigma #R8875-25G) and 0.5 μM of antimycin A (Sigma #A8674-100MG), respectively. One measuring cycle consisted of 3 minutes of mixing, 3 minutes of waiting and 3 minutes of measuring. The OCR was normalized to citrate synthase activity, to correct for the mitochondrial content of the samples (Rodenburg, 2011). The citrate synthase activity was measured according to a protocol previously described (Srere, 1969), modified for Seahorse 96 wells plates. In short, after completion of OCR measurements the Seahorse medium was replaced by 0.33% Triton X-100, 10 mM Tris-HCl (pH 7.6), after which the plates were stored at −80 °C. Before measurements, the plates were thawed and 3 mM acetyl-CoA, 1 mM DTNB, and 10% Triton X-100 were added. The background conversion of DTNB was measured spectrophotometrically at 412 nm and 37 °C for 10 minutes at 1-minute intervals, using a Tecan Spark spectrophotometer. Subsequently, the reaction was started by adding 10 mM of the citrate synthase substrate oxaloacetate, after which the ΔA_412 nm_ was measured again for 10 minutes at 1-minute intervals. The citrate synthase activity was calculated from the rate of DTNB conversion in the presence of oxaloacetate, subtracted by the background DTNB conversion rate, using an extinction coefficient of 0.0136 μmol-1. cm-1.

### Immunohistochemistry

INeurons were briefly washed with ice-cold DPBS (Gibco #14190-094), before fixation with 4% paraformaldehyde/ 4% sucrose (v/v), and subsequently permeabilized (DPBS, 0.2% triton X-100 (Sigma-Aldrich #9002-93-1)). Cells were again washed 3 times with DPBS for 5 minutes at room temperature, before a-specific binding of antibodies was prevented by incubation with blocking buffer (DPBS, 5% normal horse serum, 5% normal goat serum, 5% normal donkey serum, 0,1% bovine serum albumine (BSA), 1% glycine, 0,4% triton, 0,1% lysine (all from Sigma-Aldrich)) for 1 hour at room temperature. Primary antibodies were diluted 1:1000 in blocking buffer, and were incubated overnight, at 4°C. The cells were afterwards washed 3 times for 5 minutes at room temperature, followed by incubation with secondary antibodies, diluted 1:2000 in blocking buffer, for 1 hour at room temperature. After washing for 3 times for 5 minutes at RT with DPBS, Höchst (Termofischer #H3570) diluted 1:10.000 in DPBS was incubated for 10 minutes at RT. After one last wash with DPBS for 5 minutes at RT the coverslips were imbedded in fluorescent mounting medium (DAKO #S3023). Primary antibodies that were used, are: mouse anti-MAP2 (1:1000; Sigma M4403), and guinea pig anti-synapsin 1/2 (1:1000; Synaptic Systems 106004). Secondary antibodies that were used, are: goat anti-mouse Alexa Fluor 488 (1:2000, Invitrogen A-11029), and goat anti-guinea pig Alexa Fluor 568 (1:2000, Invitrogen). Imaging was done on a Zeiss Axio Imager Z1 equipped with apotome, using the same settings for all batches and groups, at a resolution of 1024 x 1024 at 63X magnification. Images were analyzed using ImageJ software (Schneider et al., 2012), whereby iNeurons were selected based on both Map2- and Synapsin expression. Synapsyn-puncta were counted manually and normalized to the length of the dendritic branch on which they reside.

### Neuronal morphometrical reconstruction

To examine morphology of iNeurons, cells on coverslips were transfected with plasmids expressing Discosoma species red (dsRED) fluorescent protein one week after plating. DNAin (MTI-GlobalStem) was used according to manual instructions. Medium was refreshed completely the day after DNAin application. After three weeks of differentiation cells were fixed in 4% paraformaldehyde/ 4% sucrose (v/v) in PBS and mounted with DAKO. Transfected iNeurons were imaged using a Zeiss Axio Imager Z1 and digitally reconstructed using Neurolucida 360 software (Version 11, MBF–Bioscience, Williston, ND, USA). For large cells multiple 2-dimensional images of these iNeurons were taken and subsequently stitched together using the stitching plugin of FIJI 2017 software. The 3-dimensional reconstructions and quantitative morphometrical analyses focused on the somatodendritic organization of the iNeurons. We defined origins for individual primary dendrites by identifying emerging neurites with diameters that were less than 50% of the diameter of the soma. Axons, which were excluded from reconstructions and further analysis, we visually identified by their long, thin properties, far reaching projections and numerous directional changes iNeurons that had at least two primary dendrites and reached at least the second branching order were considered for analysis. For morphometrical analysis we determined soma size, number of primary dendrites, length and branching points per primary dendrite and total dendritic length. To measure the total span of the dendritic field (receptive area) of a neuron we performed convex hull 3D analysis. Note, that due to the 2-dimensional nature of the imaging data, we collapsed the convex hull 3D data to 2-dimensions, resulting in a measurement of the receptive area and not the volume of the span of the dendritic field. Furthermore, we performed Sholl analysis to investigate dendritic complexity in dependence from distance to soma. For each distance interval (10 μm each) the number of intersections (the number of dendrites that cross each concentric circle), number of nodes and total dendritic length was measured.

### Whole-cell patch clamp recordings

Experiments were performed in a recording chamber on the stage of an Olympus BX51WI upright microscope (Olympus Life Science) equipped with infrared differential interference contrast optics, an Olympus LUMPlanFL N 40x water-immersion objective (Olympus Life Science) and kappa MXC 200 camera system (Kappa optronics GmbH) for visualisation. Through the recording chamber a continuous flow of carbongenated artificial cerebrospinal fluid (ACSF) containing (in mM) 124 NaCl, 1.25 NaH2PO4, 3 KCl, 26 NaHCO3, 11 Glucose, 2 CaCl2, 1 MgCl2 (adjusted to pH 7.4), warmed to 30°C, was present. Patch pipettes (6-8 MΩ) were pulled from borosilicate glass with filament and fire-polished ends (Science Products GmbH) using the PMP-102 micropipette puller (MicroData Instrument). For recordings of spontaneous excitatory postsynaptic currents (sEPSCs) in voltage clamp mode pipettes were filled with a cesium-based solution containing (in mM) 115 CsMeSO3, 20 CsCl, 10 HEPES, 2.5 MgCl2, 4 Na2-ATP, 0.4 Na3-ATP, 10 Na-phosphocreatine, 0.6 EGTA (adjusted to pH 7.2 and osmolarity 304 mOsmol). Recordings were made using a Multiclamp 700B amplifier (Molecular Devices, Wokingham, United Kingdom)). SEPSCs were analysed using MiniAnalysis (Synaptosoft Inc). Spontaneous action potential-evoked postsynaptic currents (sEPSC) were recorded in ACSF without additional drugs at a holding potential of −60 mV.

### Micro-electrode array recordings

Recordings of the spontaneous activity of neuronal networks derived from nine iPS lines (CTR, LH1-4, HH1-4) were performed at DIV 28. All recordings were performed using the 24-well MEA system (Multichannel Systems, MCS GmbH, Reutlingen, Germany). MEA devices allow for non-invasive recording of neuronal activity, simultaneously at multiple nodes of the same network. Each of the 24 independent wells is embedded with 12 microelectrodes (i.e. 12 electrodes/well, 30 μm in diameter and spaced 300 μm apart). The neuronal networks could acclimatize to the recording environment (37°C; 5% CO2) for 10 minutes, after which the spontaneous neuronal network activity was recorded for a subsequent 10 minutes. The signal was sampled at 10 KHz, filtered with a high-pass filter (i.e. Butterworth, 100 Hz cut-off frequency) and the noise threshold was set at ± 4.5 standard deviations. Data analysis was performed off-line by using Multiwell Analyzer (i.e. software from the 24-well MEA system that allows the extraction of the spike trains) and a custom software package named SPYCODE developed in MATLAB (The Mathworks, Natick, MA, USA) that allows the extraction of parameters describing the network activity (Bologna et al., 2010). The mean firing rate (MFR) of the network was obtained by computing the firing rate of each channel averaged among all the active electrodes of the MEA. Burst detection. Burst were at least 4 spikes in burst with a minimal inter-spike-interval of 100 milliseconds. Network burst detection. Synchronous events were detected looking for sequences of closely spaced single-channels bursts. A network burst was identified if it involves at least the 80% of the network active channels (i.e. firing rate higher that 0.1 Hz).

### Mitochondrial morphology

Analysis of mitochondrial morphology was done on transfected iNeurons using the DNA-In Neuro Transfection Reagent (Globalstem Inc), in combination with a DsRed2-Mito7 plasmid, which was a gift from Michael Davidson (Addgene plasmid # 55838). Normal differentiation protocol was followed as previously described, up to DIV6, where 100% of the medium was replaced with Neurobasal medium supplemented with 20 μg/mL B-27, 10 μg/mL glutaMAX, 10 ng/mL human recombinant NT3, 10 ng/mL human recombinant BDNF. A mix was made containing 0.5 μg DNA, 3 μl DNA-In Neuro reagents, and 22 μl Neurobasal medium, which was incubated at room temperature for 15 minutes, before being added to the wells in question. After 24-hour incubation, the medium is completely removed, and replaced with normal Neurobasal medium supplemented with 20 μg/mL B-27, 10 μg/mL glutaMAX, 1% Pen/Strep, 4 μg/mL Doxycycline, 10 ng/mL human recombinant NT3, 10 ng/mL human recombinant BDNF, after which the normal differentiation protocol is followed until DIV23. Subsequently, the cells were fixated as previously described, and an immunohistochemistry staining for MAP2 was done (see supplementary methods). Cells were imaged using a Zeiss Axio Imager Z1 equipped with apotome, using the same settings for all batches and groups at 40X magnification. Mitochondrial morphology was analysed using a specific protocol by (Valente et al., 2017), making use of the ImageJ software (Schneider et al., 2012). Morphology was analysed using a method and ImageJ plugin previously described (Dagda et al., 2009).

### Statistical analysis

The data is expressed as the mean + standard error of the mean (SEM). Analysis was done using unpaired t-tests, one-way analysis of variance with Bonferroni post hoc correction, or one-way repeated measures ANOVA with sequential post hoc Bonferroni corrections, or Kolmogorov-Smirnov test, where appropriate, using GraphPad Prism 6 (GraphPad Software). P-values of P<0.05 and smaller, were deemed significant. Sample sizes were based on our previous experiences in the calculation of experimental variability, and are described, per experiment, in the corresponding figure legends. The output of all analyses is grouped per figure and combined in supplementary table S1.

### Data availability

The authors confirm that the data supporting the findings of this study are available within the article [and/or] its’ supplementary material. Raw data supporting the presented findings of this study are available from the corresponding author upon reasonable request.

