## Supplemental data for "Mitochondrial dysfunction impairs human neuronal development and reduces neuronal network activity and synchronicity"

**Supplementary material**

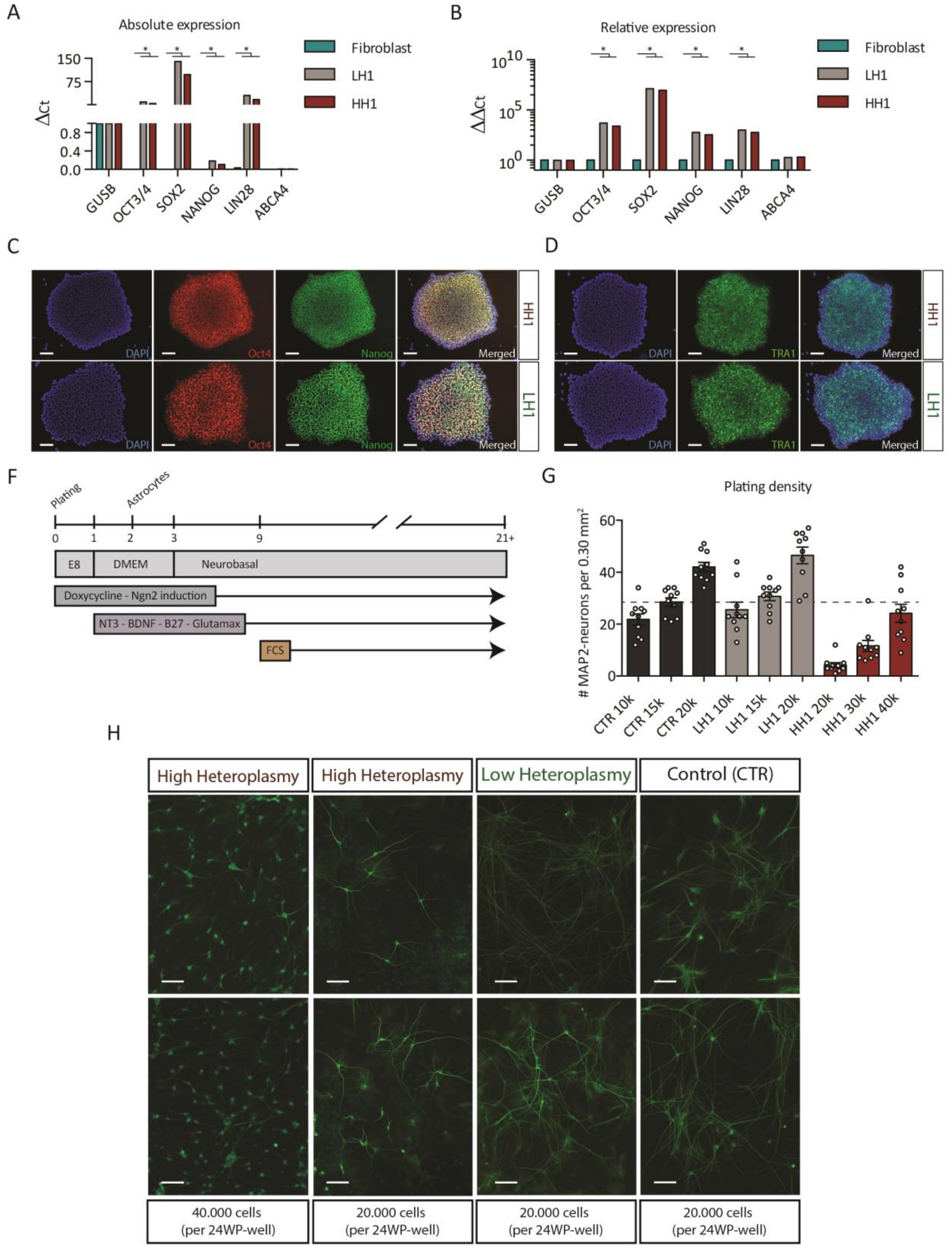


*Supplementary figure 1. Pluripotency quantification by quantitative PCR as well as immunohistochemical stainings, and neuronal density upon maturation.* (A) IPS cell pluripotency tested by quantitative real-time PCR, based on delta-ct levels for octamer-binding transcription factor 3/4 (OCT3/4), SRY-box 2 (SOX2), NANOG, and LIN28, using glucuronidase beta (GUSB) as housekeeping gene, and ABCA4 as negative control, as well as (B) the relative gene expression normalized to GUSB level. (C) Light fluorescence images of pluripotency genes OCT4, NANOG, and DAPI (nuclear marker), for LH1 and HH1 IPS lines (Scale bar = 100 µm). (D) Light fluorescence images of pluripotency gene TRA1 for LH1 and HH1 IPS lines (Scale bar = 100 µm). (F) The IPS-to-iNeuron differentiation protocol; IPS cells are plated as single cells at DIV0 supplemented with Doxycycline to induce differentiation. DIV1: Medium switch to DMEM supplemented with Doxycycline, NT3, BDNF, B27, and Glutamax. DIV2: Astrocytes are plated at 1:1 ratio. DIV3: Medium switch to Neurobasal (supplemented Doxycycline, NT3, BDNF, B27, Glutamax); DIV9: Addition of FCS to medium. (G) Neuronal density at maturity (DIV21) based on 10.000 / 15.000 / 20.000 / 30.000 / 40.000 initial plating density; quantified the number of MAP2 positive cells per 0.30 $mm^2$. (H) Light fluorescence images of MAP2 positive LH- and HH- iNeuron cultures used to quantify plating density by counting the number of MAP2 positive cells (Scale bar = 100 µm). Data represents means ± SEM, *P<0.05, **P<0.01, ***P<0.001, ****P<0.0001, one-way-, or two-way, analysis of variance with post hoc Bonferroni correction.

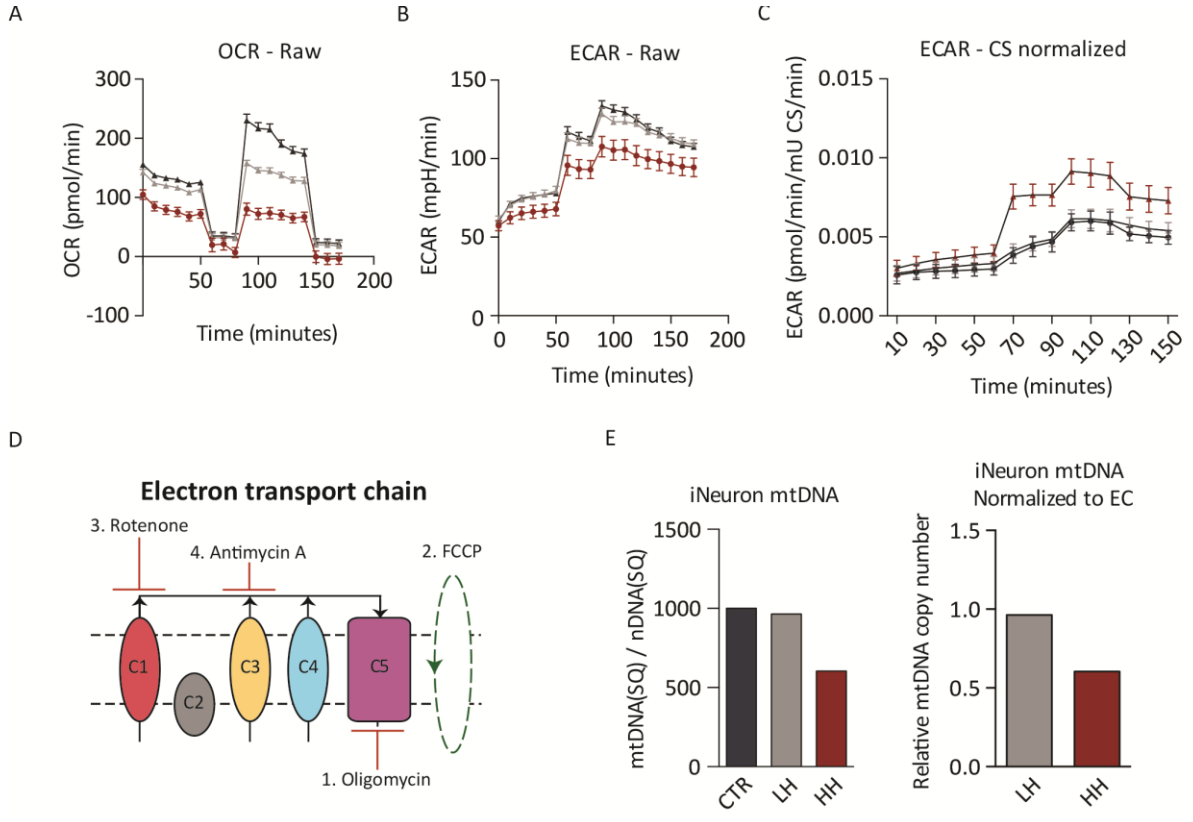


*Supplementary figure 2. Neuronal aerobic metabolic profiles of CTR, LH, and HH iNeurons as measured by Seahorse Extracellular Flux Analyzer*. (A) Raw results of fifteen consecutive oxygen consumption rate (OCR) measurements, at basal level, and following supplementation with Oligomycin (2 µM), FCCP (2 µM), and Rotenone and Antimycin A (0.5 µM). (B) Raw extracellular acidification rate (ECAR) (represents glycolysis rate) determined during- and averaged over- the first six recordings (n = 12); (C) Raw ECAR normalized to a citrate synthase assay. Data represents means ± SEM, *P<0.05, **P<0.01, ***P<0.001, ****P<0.0001, one-way analysis of variance with post hoc Bonferroni correction. CTR-, LH- and HH iNeurons were statistically compared by One-way anova. (D) Visual representation of the effects of the Oligomycin, FCCP, Rotenone, and Antimycin A, on the separate ETC complexes. C1, complex 1 (NADH dehydrogenase); C2, complex 2 (Succinate dehydrogenase); C3, complex 3 (Coenzyme Q: cytochrome c reductase); C4, complex 4 (Cytochrome c oxidase); C5, complex 5 (ATP synthase). (E) MtDNA depletion quantitative PCR used to quantify relative mtDNA levels, using SQ (standard quality) mtDNA and SQ (standard quality) nuclear DNA levels, and plotted the data relative to the external control.

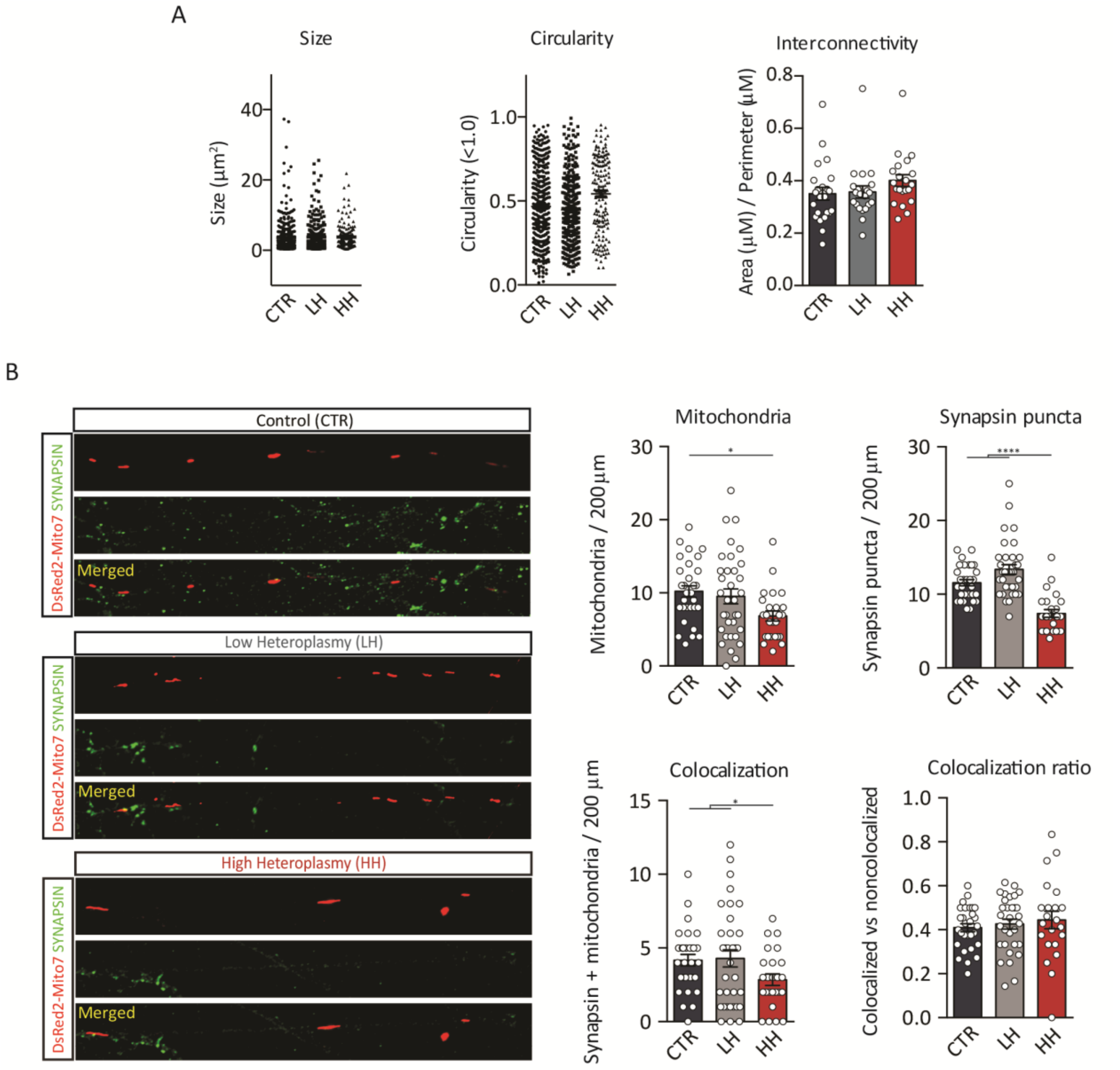


*Supplementary figure 3. Mitochondrial and synaptic density in the axon*. (A) Mitochondrial size plotted per mitochondria, for all CTR-, LH-, and HH- iNeurons analyzed (CTR *n* = 385; LH *n* = 342; HH *n* = 162). Mitochondrial circularity (1 = perfectly round) plotted per mitochondria, for all CTR-, LH-, and HH- iNeurons analyzed (CTR *n* = 385; LH *n* = 342; HH *n* = 162). Average interconnectivity (area (µm$^2$) / perimeter (µm)) plotted per iNeuron, for all CTR-, LH-, and HH- iNeurons analyzed (CTR *n* = 23; LH *n* = 21; HH *n* = 21). (B) Light fluorescence images of CTR-, LH-, and HH iNeurons transfected with a DsRed2-Mito7 (mitochondria) and stained for Synapsin1/2 (pre-synaptic protein) (Scale bar = 30 µm). Quantified the number of mitochondria, the number of Synapsin puncta, the absolute number of co-localizing (mitochondria plus Synapsin puncta) and the ratio of co-localizing (co-localizing / non co-localizing) (CTR *n* = 30; LH *n* = 31; HH *n* = 25). Data represents means ±

SEM, *P<0.05, **P<0.01, ***P<0.001, ****P<0.0001, one-way analysis of variance with Bonferroni post hoc test for multiple comparisons.

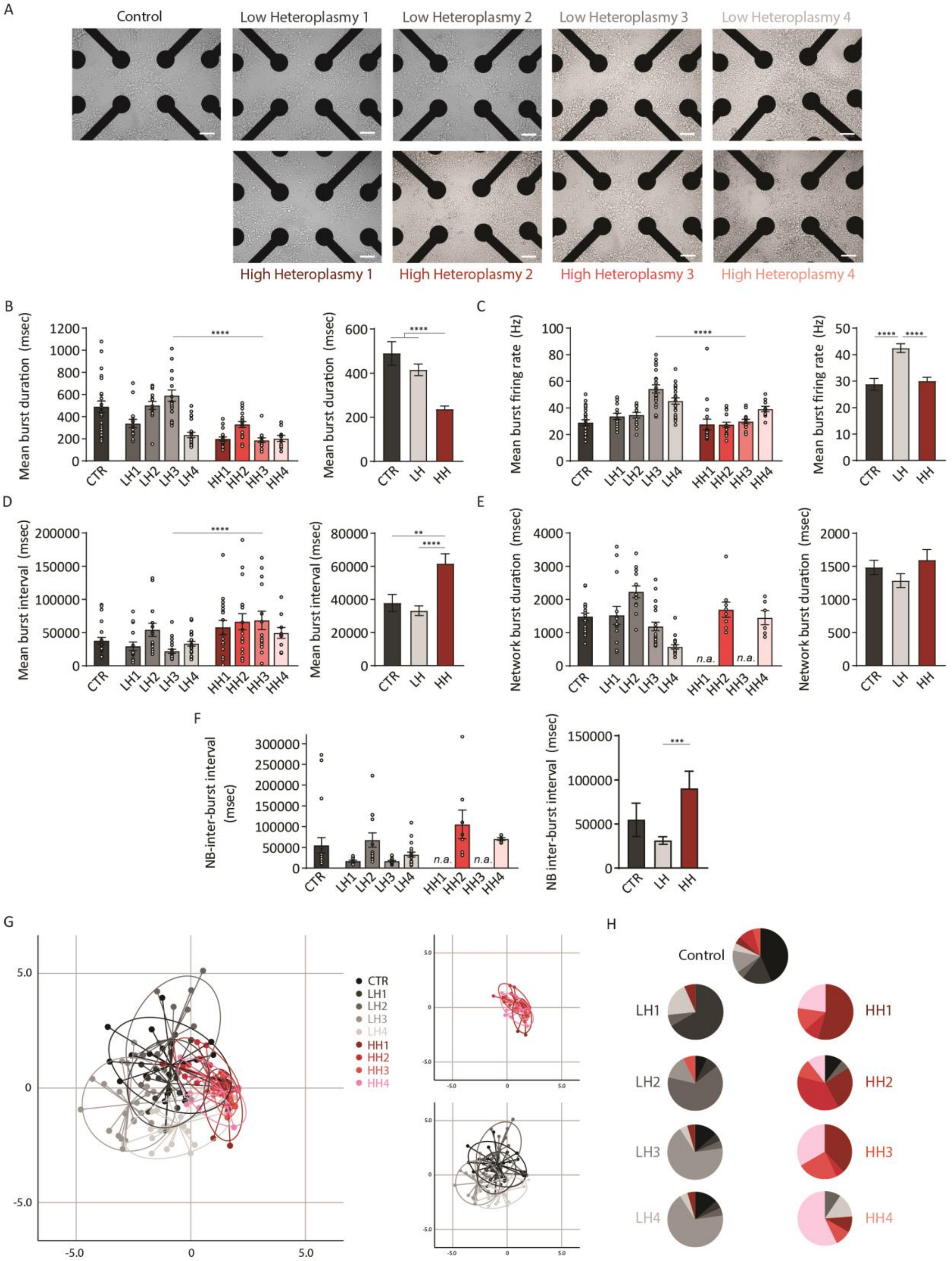


*Supplementary figure 4. CTR-, LH-, and HH Neuronal network activity.* (A) CTR, LH1-4, and HH1-4 neuronal networks were grown at similar density on 24-well micro-electrode arrays (MEA) (CTR n = 23; LH1 n = 15; LH2 = 14; LH3 = 22; LH4 = 23; HH1 n = 22; HH2 = 20; HH3 = 21; HH4 = 21), as visible in bright field images taken for CTR-, LH-, and HH MEA cultures upon maturation (Scale bar = 100 μm). We quantified (B) mean burst duration (MBD), (C) mean burst firing rate (MBFR), (D) mean burst interval (MBI), (E) network burst duration (NBR), and (F) network inter-burst interval (NIBI). Data represents means ± SEM, *P<0.05, **P<0.01, ***P<0.001, ****P<0.0001, one-way analysis of variance with Bonferroni post hoc test for multiple comparisons. (G) Unbiased discriminant function analysis with canonical discriminant functions classifies control (CTR), low heteroplasmy (LH1-4), and high heteroplasmy (HH1-4) lines based on mean firing rate (MFR), percentage of random spikes (PRS), mean burst rate (MBR), mean burst duration (MBD), mean burst firing rate (MBFR), network burst rate (NBR), network burst duration (NBD), and network inter-burst interval (NIBI). Each dot represents one experiment (CTR n = 23; LH1 n = 15; LH2 = 14; LH3 = 22; LH4 = 23; HH1 n = 22; HH2 = 20; HH3 = 21; HH4 = 21). (H) Group membership predictions by discriminant function analysis. Lines are represented by distinct colors.

| | Patients / Subjects | | | |
|---|---|---|---|---|
| **Basic information** | #1 (C1) | #2 (MitoA) | #3 (MitoC) | #4 (DB) |
| Gender | Male | *Female* | Male | *Female* |
| Origin | Leuven, Belgium | *Rochester, USA* | *Rochester, USA* | *Rochester, USA* |
| Race | Caucasian | - | - | - |
| Age (Biopsy) | 42 | 17 | 31 | 45 |
| IPSC reprogramming method | Lentiviral | *Sendai* | *Sendai* | *Sendai* |
| **M.3243A>G heteroplasmy levels** | | | | |
| Blood | 58% | 89% | - | 0-86% |
| Fibroblast | - | - | - | 0-79% |
| Urine | 28% | 23-30% | 86% | 0-84% |
| IPSC | 16% | 33% | 47% | 0-87% |
| **Symptoms** (age of onset in years) | | | | |
| Stroke-like episodes | 33-42+ | 36+ | 30+ | 30+ |
| Epilepsy / seizures | | 36+ | 30+ | |
| Recurrent headaches / migraine | | | 30+ | |
| (Progressive) hearing impairment | 33+ | 6-12+ | 25+ | 25+ |
| Cortical vision loss / optic disorders | | | 30+ | |
| Diabetes | 31+ | 12-16+ | 18+ | 23+ |
| Ragged red fibers | 42+ | | | |
| Exercise intolerance | 12+ | | | 37+ |
| Muscle weakness / rhabdomyolysis | 42+ | | | 37+ |
| Recurrent vomiting | | | 6-12+ | |
| Ataxia | | 36+ | | |
| Neuropathy / Encephalomyopathy | 42+ | | | 37+ |
| Cardiomyopathy | 31+ | | | |
| Nephropathy | | | | 25+ |
| microalbuminuria | | | | 25+ |
| Opthalmoplegia | | | | 37+ |
| Memory loss | | | | 35+ |
| Cognitive decline | | 36+ | 30+ | 35+ |
| Depression / Anxiety | 33+ | 12-16+ | 25+ | 25+ |
| Suicidal thoughts | | | | 35+ |
| Dementia | | 36+ | 30+ | |
| Loss of capacity for personal care | | | 31+ | |

*Supplementary table 1. Patient information and symptom age of onset*. The table contains information on the patient gender, city of origin, race, age (at biopsy), age (diagnosis), IPSC reprogramming method, m.3243A>G heteroplasmy levels per tissue type, and the type- and age of onset- of the symptoms that each patient has developed through the years.
